# Genetically Encoded Melanin as a Photostable Scattering Contrast for Whole-Brain Tomography

**DOI:** 10.64898/2026.06.28.735089

**Authors:** Peilin Gu, Chong Chen, Jian Ren

## Abstract

Large-scale brain imaging has traditionally depended on fluorescent reporters, but challenges like photobleaching and inconsistent signals hinder quantitative analysis in intact tissues. In response, we present MelaCAST (melanin-based scattering CAST imaging), a novel genetically encoded scattering method for whole-brain visualization. Using AAV to deliver tyrosinase allows for cell-type-specific melanin synthesis, providing a stable intracellular scattering contrast across the mouse brain. Combining tissue clearing with scattering tomography, MelaCAST offers non-photobleaching, high-throughput volumetric imaging of genetically targeted cell populations in intact brains. This technique establishes melanin as a genetically encoded scattering reporter and broadens whole-organ imaging capabilities beyond traditional fluorescence methods.

## l Introduction

Genetically encoded fluorescent proteins enable cell-type-specific labeling and visualization in intact biological tissues^1–3^. When combined with tissue-clearing methods and volumetric microscopy, fluorescent reporters have become widely used to map cellular populations and neural circuits across whole organs^4–6^. Despite their broad utility, fluorescence-based approaches have several limitations for large-scale imaging. First, fluorescence signals are susceptible to photobleaching and can be influenced by tissue processing, imaging conditions, and clearing protocols, complicating quantitative comparisons across large specimens and over extended imaging sessions^7–9^. In addition, the finite photon yield of fluorophores constrains acquisition speed, making high-resolution imaging of cleared tissues increasingly time-intensive as sample size increases^10^. These challenges are particularly relevant for whole-organ imaging studies requiring large cohorts or longitudinal analyses.

To address these problems, elastic scattering provides an alternative contrast mechanism that is inherently resistant to photobleaching. Recent developments in Clearing-Assisted Scattering Tomography (CAST) have enabled volumetric imaging of optically cleared tissues based on scattering contrast^11^. However, in contrast to fluorescence microscopy, scattering imaging currently lacks genetically encoded reporters for selective labeling of defined cell populations, limiting its integration with established molecular and viral-labeling strategies.

Here, we present **MelaCAST**, a genetically encoded scattering-labeling approach based on intracellular melanin biosynthesis. Through AAV-mediated expression of tyrosinase (TYR), melanin is produced in genetically targeted cell populations, generating scattering contrast that is compatible with tissue clearing and CAST imaging. Using this strategy, we achieve whole-brain labeling and volumetric imaging of neuronal and astrocytic populations in intact mouse brains. This work establishes melanin as a genetically encoded scattering reporter and provides a framework for combining genetic targeting with large-scale scattering-based imaging.

## 2. Result and discussion

### 2.1 Genetically encoded melanin production enables scattering-based imaging

To establish a genetically encoded scattering reporter for CAST imaging, we developed AAV vectors expressing TYR, the key enzyme responsible for melanin biosynthesis^12^ (Fig. 1a,b).

**Figure 1.**
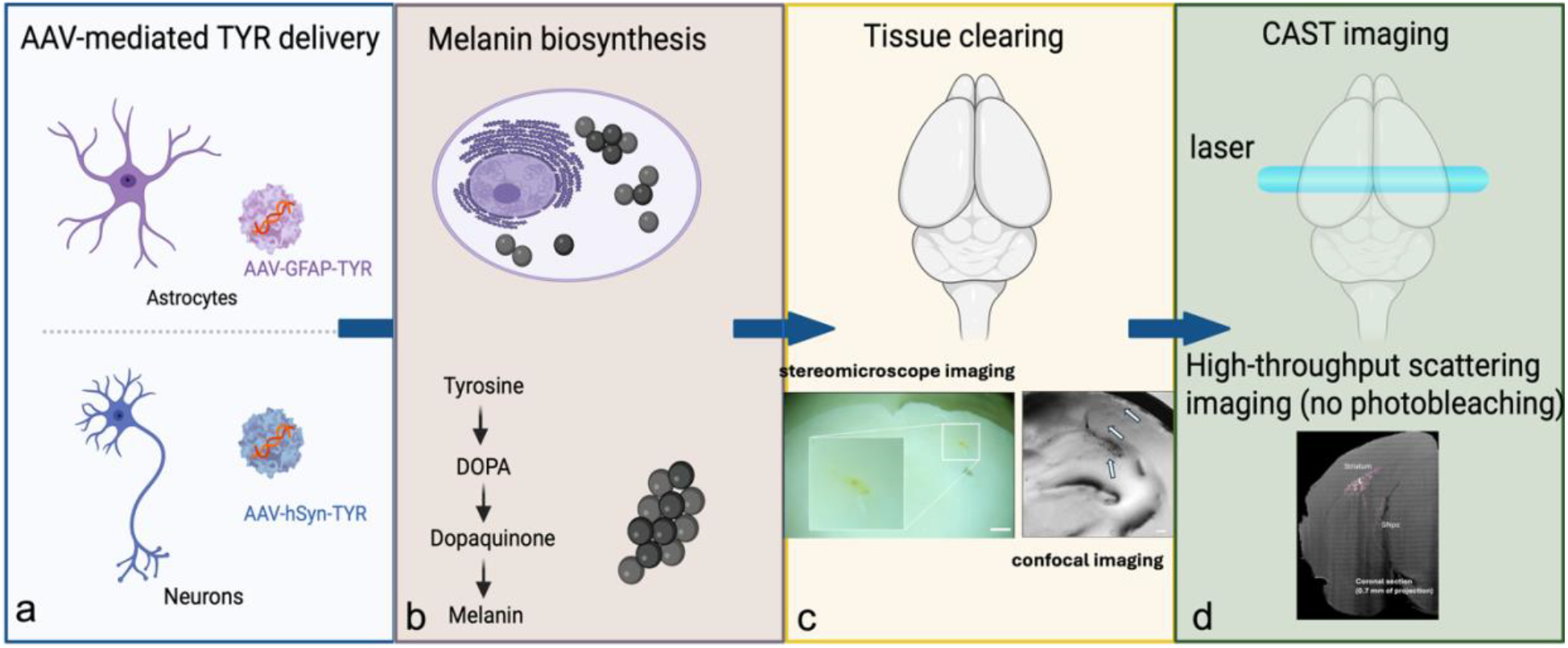
Establishment of a genetically encoded melanin reporter for CAST imaging. **a**, Schematic illustration of the AAV-TYR construct used for genetically encoded melanin labeling. **b**, Mechanism of TYR-mediated melanin biosynthesis. Following AAV transduction, intracellular expression of TYR catalyzes the conversion of tyrosine into melanin, generating endogenous optical scattering contrast. **c**, Validation of melanin production in vivo following intracerebroventricular injection of AAV2-EF1α-TYR. Stereomicroscope imaging shows visible dark pigmentation at the injection site in the mouse brain. After tissue clearing, confocal imaging of the same region reveals abundant intracellular melanin deposition. **d**, CAST imaging of adjacent brain sections. Strong scattering signals were detected in regions containing melanin deposits, showing close spatial correspondence between melanin accumulation and CAST contrast.

Following viral transduction, intracellular TYR expression catalyzes the conversion of tyrosine into melanin, generating endogenous scattering contrast within genetically targeted cells. As an initial proof of concept, AAV2-EF1α-TYR was administered by intracerebroventricular injection to induce local melanin production in the mouse brain. Eight weeks after injection, brain tissues were harvested and examined using conventional optical imaging methods. First, stereomicroscope imaging revealed visible dark pigmentation around the injection site (Fig. 1c), indicating successful melanin production in vivo. After tissue clearing, confocal imaging of the same region further confirmed the presence of abundant melanin deposits within the transduced area (Fig. 1c). We next performed CAST imaging on adjacent tissue sections. Strong scattering signals were detected in the same brain regions identified by bright-field imaging, demonstrating a close correspondence between melanin deposition and CAST contrast (Fig. 1d). Together, these results demonstrate that AAV-mediated TYR expression successfully induces melanin production in targeted brain regions and that the resulting melanin generates strong scattering contrast detectable by CAST. These findings establish the feasibility of using genetically encoded melanin as a scattering reporter for subsequent cell-type-specific whole-brain imaging.

### 2.2 Melanin labeling exhibits high astrocyte specificity and minimal overt toxicity

To evaluate the specificity and efficiency of TYR-mediated melanin labeling, AAV.PHP.eB-GFAP-TYR was administered via retro-orbital injection to induce astrocyte-targeted melanin production throughout the brain. Following an eight-week expression period, brain tissues were collected and sectioned for histological analysis. To validate the cellular identity of melanin-labeled cells, brain sections were subjected to Masson-Fontana staining in combination with immunofluorescence labeling of astrocytes^13^. Masson-Fontana-positive melanin deposits showed extensive colocalization with GFAP-positive astrocytes in the hippocampus and cortex (Fig. 2a,b), indicating preferential melanin production in astrocytes. We next quantified the labeling efficiency and labeling specificity of the GFAP-driven TYR construct. In the hippocampus, the transduction efficiency reached 86.5%, whereas in the cortex it reached 93.1% (Fig. 2c). The specificity of melanin labeling for astrocytes was 90.3% in the hippocampus and 89.0% in the cortex (Fig. 2d). These values are comparable to previously reported transduction efficiencies and cell-type specificities achieved using AAV.PHP.eB-mediated fluorescent protein expression, demonstrating that TYR-mediated melanin production enables efficient and selective astrocyte labeling throughout the brain^14,15^.

**Figure 2.**
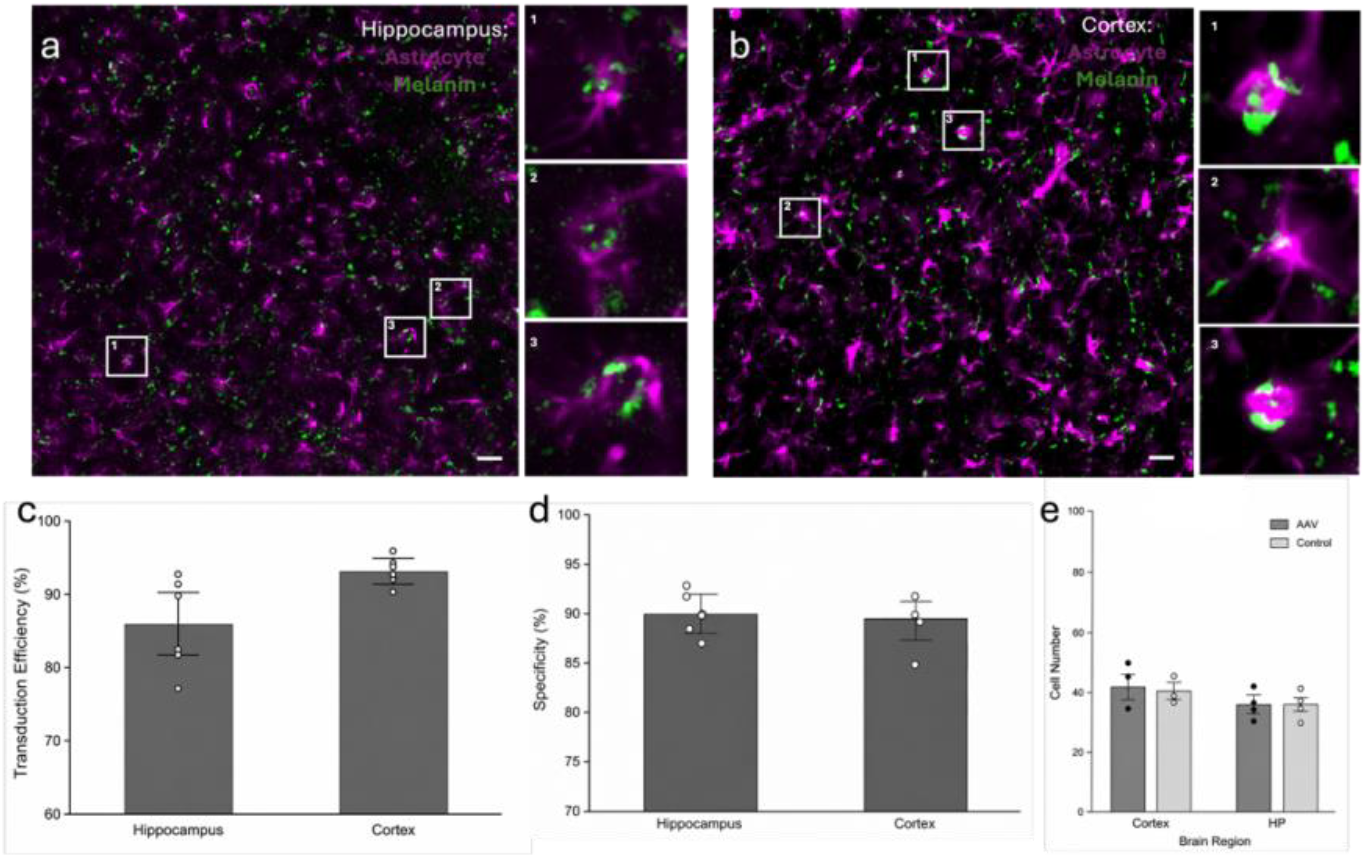
Astrocyte-specific melanin labeling following systemic delivery of AAV.PHP.eB-GFAP-TYR. **a, b**, Representative images of Masson-Fontana staining (green) and GFAP immunofluorescence (magenta) in the hippocampus and cortex. **c**, Quantification of transduction efficiency in the hippocampus and cortex. **d**, Quantification of labeling specificity in the hippocampus and cortex. **e**, Representative images of astrocyte staining and quantification of astrocyte density in control and TYR-expressing mice. Data are presented as mean ± s.d. Scale bars, 10 μm.

To assess the biosafety of intracellular melanin production, brain sections from TYR-expressing and control mice were immunostained for astrocytic markers, and cell densities were quantified across multiple brain regions. No significant differences in cell number or gross cellular morphology were observed between groups (Fig. 2e), suggesting that TYR-mediated melanin production does not induce overt structural toxicity under the conditions examined.

Together, these results demonstrate that AAV-mediated TYR expression enables efficient and astrocyte-specific melanin labeling while exhibiting minimal detectable neurotoxicity, supporting its application as a genetically encoded reporter for large-scale brain imaging.

### 2.3 CAST signals originate from genetically encoded melanin

To validate the origin of the CAST signal, 50-μm-thick cleared brain sections were first imaged by CAST and subsequently stained using the Masson-Fontana method for melanin detection. Brightfield microscopy revealed melanin deposits that closely matched the spatial distribution of CAST signals (Fig. 3). Strong colocalization between melanin staining and CAST contrast confirmed that the observed scattering signals originated from intracellular melanin rather than endogenous tissue structures. These results demonstrate that genetically encoded melanin remains detectable following tissue clearing and generates robust scattering contrast for CAST imaging, supporting its use as a reporter for large-scale volumetric imaging of genetically defined cell populations.

**Figure 3.**
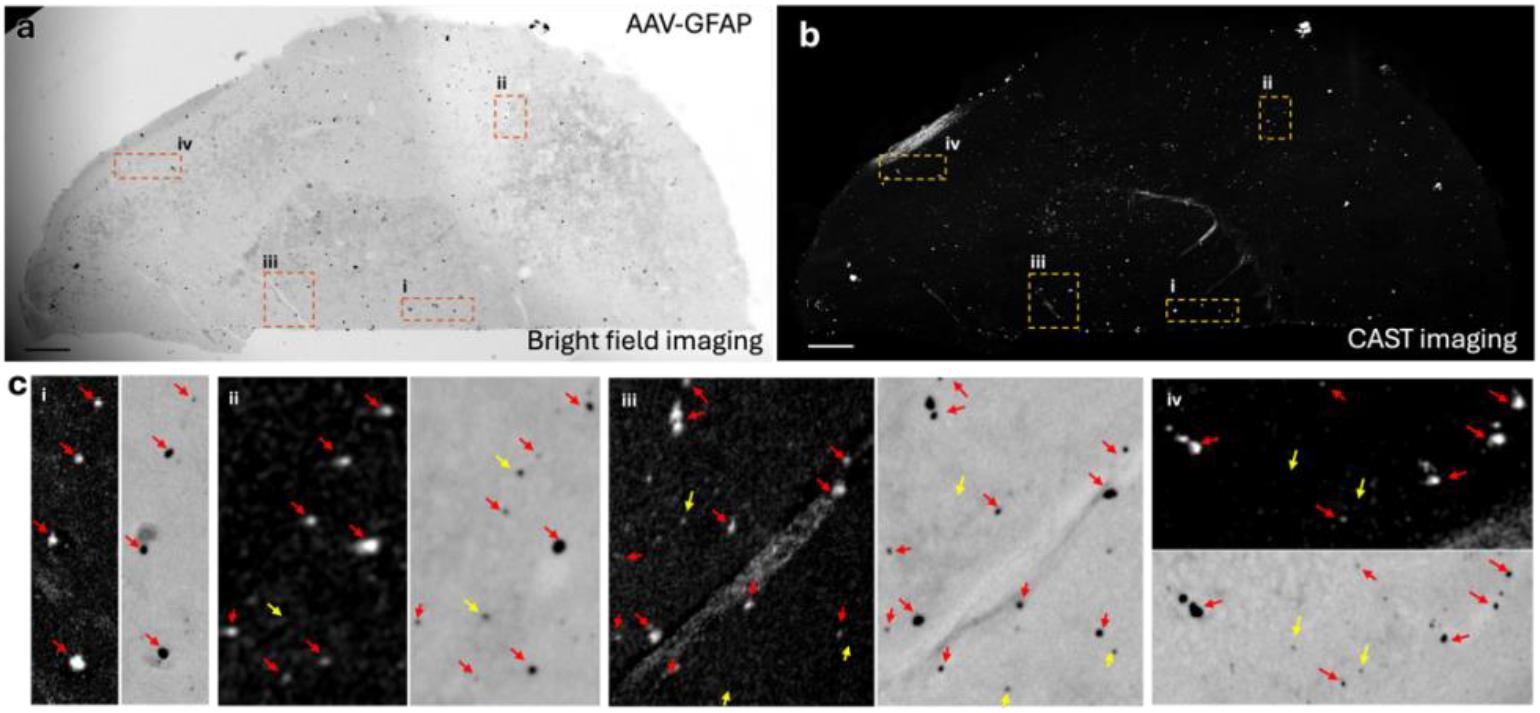
Comparison of CAST imaging and Masson-Fontana staining in melanin-labeled brain sections. **a**, Bright-field image of a brain section following Masson-Fontana staining. Dashed boxes indicate regions shown at higher magnification in c. **b**, CAST image of the same brain section acquired prior to Masson-Fontana staining. Dashed boxes indicate regions shown in c. **c**, Higher-magnification views of the regions indicated in a and b. Corresponding CAST images (left) and bright-field images after Masson-Fontana staining (right) are shown for each region (i–iv). Red arrows indicate structures detected in both CAST and Masson-Fontana images, whereas yellow arrows indicate structures detected in only one modality. Scale bars, 500 μm in a,b.

### 2.4 Whole-brain CAST imaging of genetically defined cell populations

Having established and validated the cell-type-specific scattering labeling strategy, we next evaluated its generalizability across different cellular populations. In addition to the astrocyte-targeting construct (AAV.PHP.eB-GFAP-TYR), we generated two additional AAV vectors: AAV.PHP.eB-hSyn-TYR for pan-neuronal labeling and AAV.PHP.eB-mTH-TYR for selective labeling of dopaminergic neurons^16^.

All three vectors were administered via retro-orbital injection and allowed to express for 8 weeks. Following tissue collection and optical clearing, whole brains were imaged using CAST (Fig. 4). Robust melanin-derived scattering contrast was observed throughout the brain in all three labeling paradigms, demonstrating the compatibility of TYR-mediated melanin production with whole-brain CAST imaging.

**Figure 4.**
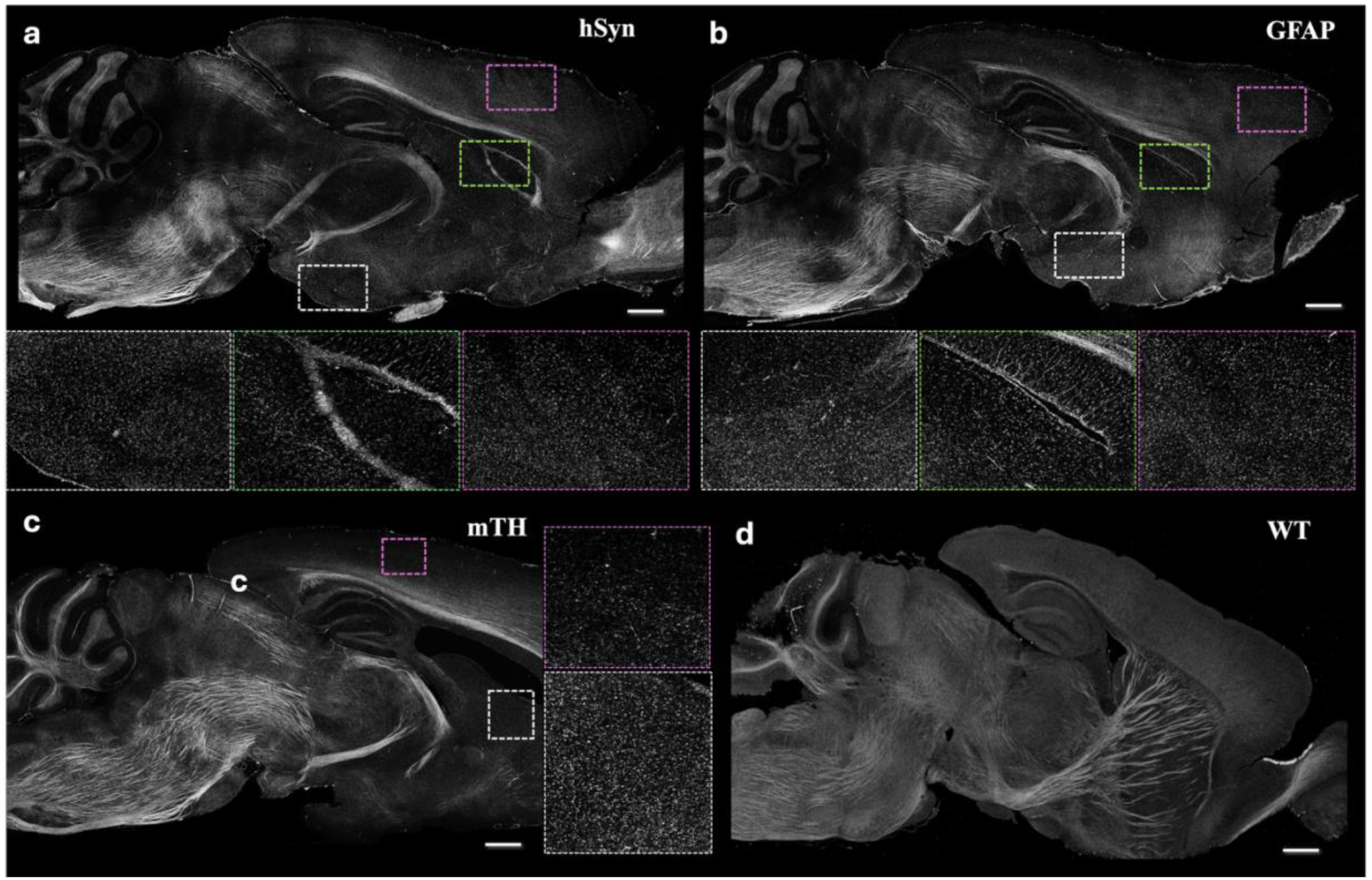
Whole-brain CAST imaging of genetically defined cell populations. **a**, Representative whole-brain CAST images of mice expressing TYR under the hSyn promoter, enabling pan-neuronal melanin labeling. Corresponding high-magnification views highlight cellular and regional labeling patterns in selected brain areas. **b**, Representative whole-brain CAST images of mice expressing TYR under the GFAP promoter, resulting in astrocyte-specific melanin labeling. Enlarged views show the distribution and morphology of labeled astrocytes in representative regions. **c**, Representative whole-brain CAST images of mice expressing TYR under the mTH promoter, labeling dopaminergic neuronal populations. Corresponding magnified images illustrate the spatial distribution of labeled cells and projections. **d**, Wild-type control mouse brain imaged under identical CAST conditions, showing the absence of exogenous melanin-derived scattering contrast. Scale bars, 500 μm.

The GFAP-driven construct enabled brain-wide visualization of astrocyte distributions, whereas hSyn-driven expression produced widespread neuronal labeling across multiple brain regions. In contrast, the mTH-driven construct selectively labeled dopaminergic neuronal populations, allowing visualization of their characteristic anatomical distributions. In all cases, melanin-generated scattering signals were readily detected by CAST, enabling volumetric reconstruction of genetically defined cell populations throughout the intact brain. As a negative control, a wild-type mouse that did not receive TYR transduction was imaged under identical conditions. No cell-associated scattering signals were detected, confirming that the observed CAST contrast originated from TYR-induced melanin production.

Together, these results demonstrate that MelaCAST can be readily adapted to different promoters and cell types, establishing a versatile platform for genetically targeted, whole-brain scattering imaging and large-scale cellular mapping.

## 3. Conclusion

In this study, we developed MelaCAST, a genetically encoded scattering-labeling strategy that combines AAV-mediated tyrosinase expression, intracellular melanin biosynthesis, tissue clearing, and CAST imaging. Using this approach, we achieved cell-type-specific melanin labeling and whole-brain scattering imaging of astrocytes, neurons, and dopaminergic neurons. These results establish melanin as a genetically encoded scattering reporter and extend genetic labeling strategies beyond fluorescence-based imaging.

A major advantage of MelaCAST is its compatibility with large-scale volumetric imaging. Unlike fluorescent reporters, melanin generates contrast through elastic scattering and is therefore resistant to photobleaching. In addition, melanin-derived contrast is preserved following tissue clearing and can be readily detected by CAST, enabling high-throughput imaging of intact brains. By combining the targeting flexibility of AAV vectors with scattering-based imaging,MelaCAST provides a practical approach for mapping genetically defined cell populations across large tissue volumes.

Several limitations should be noted. Although no overt structural toxicity was observed, the long-term effects of intracellular melanin production require further investigation^17^. In addition, the current implementation provides a single scattering contrast and does not yet offer the multiplexing capability of fluorescence imaging. Future development of additional scattering reporters and contrast-encoding strategies may further expand the utility of this approach. Overall, MelaCAST provides a framework for integrating genetic targeting with scattering-based imaging and offers a complementary alternative to fluorescence for whole-brain and whole-organ imaging.

## Acknowledgements

This research was funded by National Institutes of Health (NIH) (R00AG059946)

## Methods

### Plasmid construction

All AAV plasmids were designed and constructed by WZ Biosciences (WZ Biosciences, USA). Construct identity and sequence verification were performed by the provider prior to viral packaging.

### AAV production and stereotaxic delivery

Recombinant AAVs were produced by the Viral Vector Core Facility at Boston Children’s Hospital (BCH). Viral preparations were used without further purification. For intracerebroventricular (ICV) delivery, AAV2 was injected into adult mice at a final dose of 1 μL per animal with a viral titer of 2 × 10^13^ vg/mL. For systemic delivery, AAV-PHP.eB was administered via retro-orbital injection at a dose of 1 × 10^12^ viral genomes (vg) per mouse. All injections were performed under appropriate anesthesia using standard procedures.

### Tissue fixation and sectioning

Mice were transcardially perfused with phosphate-buffered saline (PBS), followed by fixation with 4% paraformaldehyde (PFA) and 1% glutaraldehyde (GA). Brains were post-fixed in the same solution overnight at 4°C. Fixed tissues were subsequently processed and sectioned using a vibratome to obtain free-floating sections for histological analysis.

### Immunofluorescence and histological staining

Standard immunofluorescence staining procedures were performed on free-floating brain sections. Briefly, sections were permeabilized, blocked, and incubated with primary antibodies followed by fluorescent secondary antibodies according to conventional protocols. Melanin visualization was performed using a Masson–Fontana staining kit following the manufacturer’s instructions. Stained sections were imaged using an Olympus fluorescence microscope under appropriate excitation/emission settings.

### Whole-brain clearing and refractive index matching

For whole-brain imaging, dissected brains were fixed in 4% PFA and 1% GA and subsequently subjected to tissue clearing. Samples were incubated in 8% SDS solution at 37°C for 3 weeks with gentle agitation until sufficient optical transparency was achieved. After clearing, brains were thoroughly washed and incubated in a 1.46 refractive index-matching solution prior to imaging to minimize light scattering^18^.

### CAST imaging

Cleared and index-matched whole brains were imaged using CAST. Imaging was performed on intact specimens under standardized acquisition settings to obtain volumetric scattering contrast throughout the entire brain.

Image processing and data analysis. Raw imaging data were processed using ImageJ (NIH). Standard preprocessing steps included contrast adjustment, background subtraction, and region-of-interest (ROI) quantification when applicable. No additional proprietary software was used.

## Reference

1. Tsien, R. Y. Constructing and Exploiting the Fluorescent Protein Paintbox (Nobel Lecture). Angewandte Chemie International Edition 48, 5612–5626 (2009).

2. Hunker, A. C. et al. Enhancer AAV toolbox for accessing and perturbing striatal cell types and circuits. Neuron 113, 1507–1524.e17 (2025).

3. Hunker, A. C. et al. Technical and biological sources of noise confound multiplexed enhancer AAV screening. Nature Communications 17, 3738 (2026).

4. Bonnavion, P. et al. Striatal projection neurons coexpressing dopamine D1 and D2 receptors modulate the motor function of D1- and D2-SPNs. Nat Neurosci 27, 1783–1793 (2024).

5. Gao, Y. et al. VIVIT: Resolving trans-scale volumetric biological architectures via ionic glassy tissue. Cell S009286742500813X (2025) doi:10.1016/j.cell.2025.07.023.

6. Glaser, A. et al. Expansion-assisted selective plane illumination microscopy for nanoscale imaging of centimeter-scale tissues.

7. Ueda, H. R. et al. Tissue clearing and its applications in neuroscience. Nat Rev Neurosci 21, 61–79 (2020).

8. Feng, R., Xie, J. & Gao, L. EDTP enhances and protects the fluorescent signal of GFP in cleared and expanded tissues. Sci Rep 14, 15279 (2024).

9. Park, Y.-G. et al. Protection of tissue physicochemical properties using polyfunctional crosslinkers. Nat Biotechnol 37, 73–83 (2019).

10. Xu, F. et al. High-throughput mapping of a whole rhesus monkey brain at micrometer resolution. Nature Biotechnology 39, 1521–1528 (2021).

11. Ren, J., Choi, H., Chung, K. & Bouma, B. E. Label-free volumetric optical imaging of intact murine brains. Scientific Reports 7, 46306 (2017).

12. Gargiulo, A. et al. AAV-mediated Tyrosinase Gene Transfer Restores Melanogenesis and Retinal Function in a Model of Oculo-cutaneous Albinism Type I (OCA1). Mol Ther 17, 1347–1354 (2009).

13. Carballo-Carbajal, I. et al. Brain tyrosinase overexpression implicates age-dependent neuromelanin production in Parkinson’s disease pathogenesis. Nat Commun 10, 973 (2019).

14. Deverman, B. E. et al. Cre-dependent selection yields AAV variants for widespread gene transfer to the adult brain. Nature Biotechnology 34, 204–209 (2016).

15. Chan, K. Y. et al. Engineered AAVs for efficient noninvasive gene delivery to the central and peripheral nervous systems. Nat Neurosci 20, 1172–1179 (2017).

16. Wang, M. et al. Whole-brain 3D imaging of dopaminergic neurons and glial cells in the mouse model of Parkinson’s disease induced by 6-OHDA. Frontiers in Aging Neuroscience Volume 17–2025, (2025).

17. Iannitelli, A. F. et al. Tyrosinase-induced neuromelanin accumulation triggers rapid dysregulation and degeneration of the mouse locus coeruleus. bioRxiv https://doi.org/10.1101/2023.03.07.530845 (2024) doi:10.1101/2023.03.07.530845.

18. Park, J. et al. Integrated platform for multiscale molecular imaging and phenotyping of the human brain. Science 384, eadh9979 (2024).

